# Open reading frame correction using antisense oligonucleotides for the treatment of cystic fibrosis caused by CFTR-W1282X

**DOI:** 10.1101/2021.08.11.455834

**Authors:** Wren E. Michaels, Cecilia Pena-Rasgado, Rusudan Kotaria, Robert J. Bridges, Michelle L. Hastings

**Affiliations:** Center for Genetic Diseases, Chicago Medical School, Rosalind Franklin University of Science and Medicine, North Chicago, IL, 60064, United States of America; School of Graduate and Postdoctoral Studies, Rosalind Franklin University of Science and Medicine, North Chicago, IL, 60064, United States of America

## Abstract

*CFTR* gene mutations that result in the introduction of premature termination codons (PTCs) are common in cystic fibrosis (CF). This mutation type causes a severe form of the disease, likely because of low *CFTR* mRNA expression as a result of nonsense mediated mRNA decay (NMD), as well as production of a non-functional, truncated CFTR protein. Current therapeutics for CF, which target residual protein function, are less effective in patients with these types of mutations, due in part to low CFTR protein levels. Splice-switching antisense oligonucleotides (ASOs) designed to induce skipping of exons in order to restore the mRNA open reading frame have shown therapeutic promise pre-clinically and clinically for a number of diseases. We hypothesized that ASO-mediated skipping of CFTR exon 23 would recover CFTR activity associated with terminating mutations in the exon, including *CFTR* p.W1282X, the 5^th^ most common mutation in CF. Here, we show that CFTR lacking the amino acids encoding exon 23 is partially functional and responsive to corrector and modulator drugs currently in clinical use. ASO-induced exon 23 skipping rescued CFTR expression and chloride current in primary human bronchial epithelial cells isolated from homozygote CFTR-W1282X patients. These results support the use of ASOs in treating CF patients with *CFTR* class I mutations in exon 23 that result in unstable *CFTR* mRNA and truncations of the CFTR protein.

**Significance Statement:** Frameshift and nonsense mutations pose a major problem for disease therapeutic development. Eliminating these mutations from the mRNA by inducing exon skipping is a relatively unexplored treatment approach, though it has shown promise for some diseases. Here, we show that eliminating a common stop mutation associated with cystic fibrosis by inducing skipping of the exon it is located in, results in a restoration of the open reading frame and recovers CFTR protein function in a manner expected to be therapeutic in CF patients who don’t currently have effective treatment options. These results are an important advancement for the cystic fibrosis community but also have implications for other diseases where terminating mutations are responsible for dysfunction.

## Introduction

Cystic fibrosis (CF) is an autosomal recessive genetic disease caused by mutations in the cystic fibrosis transmembrane conductance regulator (*CFTR*) gene (1). CFTR transports chloride and bicarbonate across the apical surface of epithelial cells. Loss of CFTR expression or function affects multiple organ systems, including the lungs, liver, pancreas, intestines, smooth muscle and heart (2, 3). In the lung, CFTR-mediated chloride secretion and sodium absorption by the epithelial sodium channel regulate airway surface liquid hydration. Loss of CFTR causes disruption of mucociliary clearance, resulting in the proliferation of airway pathogens, chronic infection, inflammation, and bronchial damage.

Though there are over 2,000 variants in *CFTR*, most therapeutics that are clinically available or in development are designed for specific patient subpopulations with the most common mutations (3, 4). Currently, there are four FDA-approved drugs that directly target CFTR function. These drugs are referred to as CFTR modulators. Ivacaftor (VX-770) potentiates function of CFTR by increasing the probability of channel opening for gating variants (5–7), lumacaftor (VX-809) and tezacaftor (VX-661) correct processing and trafficking of CFTR to the cell surface (8–12). A new corrector, elexacaftor (VX-445), works in combination with tezacaftor and ivacaftor (13, 14). While drug development has recently expanded drastically there is a critical need for therapies to treat patients with rare *CFTR* mutations, in particular nonsense variants that create a premature termination codons (PTC) resulting in low CFTR expression (4, 15).

One of the most common nonsense mutations associated with CF is CFTR p.W1282X (c.3846G>A). CFTR-W1282X is the fifth most common CF-causing mutation worldwide and the second most common class I mutation associated with the disease (http://cftr2.org). This mutation results in a truncated CFTR (CFTR_1281_), removing ∼60% of the nucleotide binding domain 2 (NBD2) but retaining 1281 of the 1480 amino acids in the full-length protein. The truncated CFTR-W1282X protein has processing and/or gating defects but is responsive to potentiator and corrector treatment (16–21). However, *CFTR-W1282X* mRNA is degraded by nonsense mediated mRNA decay (NMD), leading to a decrease in mRNA and protein abundance, thereby limiting the effect of modulator drugs (21, 22). Small molecule compounds that inhibit NMD have been shown to increase CFTR-W1282X expression but to date no effective drug candidates targeting NMD have been approved for use (21, 23).

Antisense oligonucleotides (ASOs) are another possible therapeutic approach for treating CF caused by nonsense and frameshift mutations. ASOs are short oligonucleotides, chemically-modified to create stable, specific and long lasting drugs that can be designed to modulate pre-mRNA splicing (24, 25). ASOs can be designed to block splicing and induce skipping of an exon, effectively removing it from the mRNA. This strategy can be useful as a potential therapeutic for *CFTR-W1282X* as amino acid 1282 resides in exon 23 which is a symmetrical exon that can be eliminated from the mRNA without disrupting the *CFTR* open reading frame. This approach would have a therapeutic benefit not only by producing a potentially functional protein isoform with a restored C-terminus, as suggested by biochemical and functional studies of CFTR (26–28), but also by eliminating the PTC, stabilizing the mRNA and increasing protein expression, thereby improving efficacy of CFTR modulators.

Here, we demonstrate that a CFTR isoform lacking the amino acids encoded by exon 23 has partial activity when exposed to CFTR modulator drugs. We identify a splice-switching ASO strategy that induces exon 23 skipping and show that ASO-mediated exon 23 skipping in *CFTR-W1282X* RNA, in both an immortalized human *CFTR-W1282X* bronchial epithelial cell line and primary epithelial cells isolated from CF patients homozygous for *CFTR-W1282X*, stabilizes the *CFTR* mRNA and recovers CFTR activity. We also provide evidence that ASO-induced exon 23 skipping has partial allele specificity for *CFTR-W1282X* which could be advantageous in treating CF patients heterozygous for *CFTR-W1282X* and another mutation less responsive to current therapeutics.

## Results

### CFTR mRNA lacking exon 23 generates a partially active protein

The *CFTR-W1282X* mutation resides within exon 23 of *CFTR* RNA. Exon 23 is a symmetrical exon that can be removed without disrupting the open reading frame (**Figure 1A**). As a first step in determining whether correction of the *CFTR-W1282X* open reading frame by removing exon 23 might be therapeutic, we tested whether an expressed CFTR isoform lacking the amino acids encoded by exon 23 had channel activity (**Figure 1B**). FRT cells, which lack endogenous *CFTR*, were transfected with plasmids expressing *CFTR* without exon 23 (CFTR-Δ23) or with the W1282X mutation (CFTR-W1282X). Transepithelial resistance measurements were recorded from the cells after monolayers formed and transepithelial conductance (Gt=1/Rt) attributed to forskolin-stimulated activation of CFTR and inhibition by CFTR inhibitor, Inh-172, were calculated (29). Measurements were plotted as conductance traces (**Figure 1C**) and the area under the curve (AUC) was calculated for comparison (**Figure 1D**). To test the responsiveness of CFTR-W1282X and CFTR-Δ23 to CFTR modulators known to increase CFTR-W1282X function, the cells were treated with VX-770 and C18 (VRT-534), an analog of the corrector VX-809, or VX-445 + VX-661 (13, 14, 30–33). CFTR-specific activity was undetectable in untreated or VX-770-treated cells. In contrast, both CFTR-Δ23 and CFTR-W1282X had similar significant increases in activity following corrector and potentiator treatment (**Figure 1C,D**). This increase in functional activity corresponded with an increase in CFTR protein as indicated by an increase in the fully glycosylated CFTR isoform (Band C) and the core glycosylated isoform (Band B) (**Figure 1E,F**) (8). These results demonstrate that CFTR-Δ23 has functional activity in the presence of CFTR modulator drugs.

**Figure 1.**
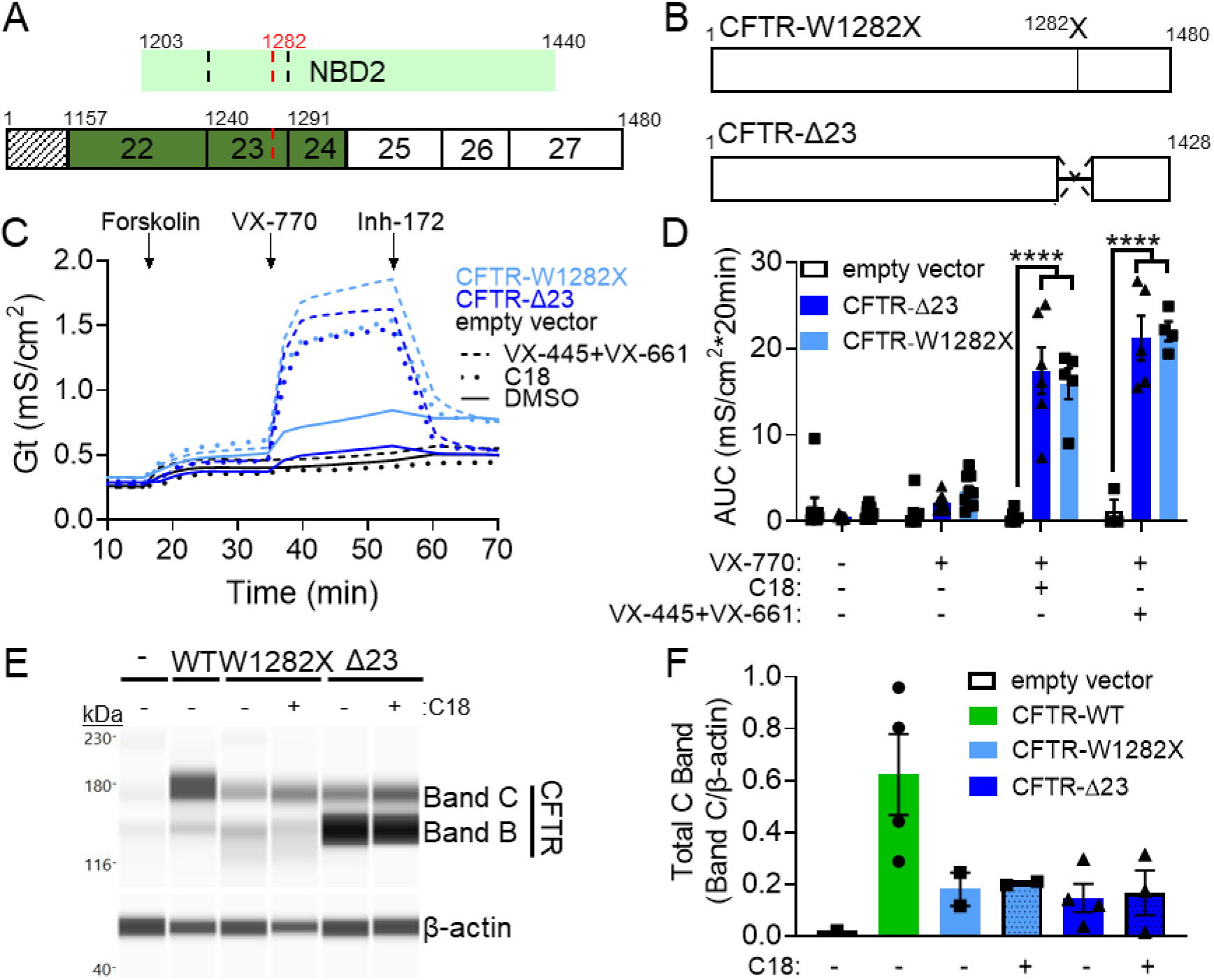
CFTR-Δ23 function is comparable to CFTR-W1282X and is responsive to CF modulators. (**A**) Schematic of CFTR exon 23 in relation to its position within NBD2 and the position of the CFTR-W1282X mutation. Symmetric exons are colored green. (**B**) Schematic of the CFTR-W1282X and CFTR-Δ23 constructs transfected into FRT cells. (**C**) Average conductance traces from FRT cells stably transfected with CFTR-Δ23, CFTR-W1282X, or empty vector. Cells were pre-treated with vehicle (DMSO, solid lines), C18 (dotted lines), or VX-445+VX-661 (dashed lines). The time of compound additions (Forskolin, VX-770, or Inh-172) is indicated. (**D**) Average area under the curve (AUC) was quantified for the forskolin and VX-770 test periods for each construct. Error bars are ±SEM. Two-way ANOVA; Dunnett’s multiple comparison test to vehicle within groups, ****p<0.0001. CFTR-Δ23: DMSO and VX-770, N=11; C18, N=6; VX-445+VX-661, N=5. CFTR-W1282X: DMSO and VX-770, N=9; C18, N=5; VX-445+VX-661, N=4. empty vector: DMSO and VX-770, N=8; C18, N=5; VX-445+VX-661, N=3. (**E**) Immunoblot analysis of CFTR protein, Bands C and B, isolated from in FRT cells stably transfected with empty vector, CFTR-WT, CFTR-W1282X, or CFTR-Δ23 constructs treated with vehicle or C18. β-actin was used as a control for protein expression. (**F**) Quantification of total CFTR C band from (E), normalized to β-actin. Error bars are ±SEM. Empty vector: N=1, CFTR-WT: N=4, CFTR-W1282X: N=2, CFTR-Δ23: DMSO, N=4; C18 N=3.

### A splice-switching ASO induces exon 23 skipping and increases *CFTR* mRNA and chloride channel activity in a *CFTR-W1282X* immortalized human bronchial epithelial cell line

Splice-switching ASOs are a therapeutic platform that can be used to induce exon 23 skipping to stabilize *CFTR-W1282X* mRNA and increase abundance of the partially functional CFTR-Δ23 protein isoform (**Figure 2A**). We tested four ASOs designed to base-pair to human *CFTR* exon 23 pre-mRNA and induce exon 23 skipping via a steric block of the splicing machinery (**Figure 2B**). ASO-23A, which basepairs to the 5’ splice site (**Figure 2B,D**), induced exon 23 skipping when transfected into an immortalized patient-derived bronchial epithelial cell line expressing *CFTR-W1282X* (CFF16HBEge-W1282X) (**Figure 2C**) (34). ASO-23A also induced splicing at a cryptic 5’ splice site within exon 23, which results in out-of-frame mRNA. To reduce the use of this cryptic splice site and maximize exon 23 skipping, cells were co-transfected with ASO-23A and another ASO, ASO-23B, which blocks the cryptic splice site (**Figure 2D**). Treatment of cells with ASO-23A and ASO-23B (ASO-23AB) eliminated cryptic splice site use and resulted in a dose-dependent increase in exon 23 skipping (**Figure 2E,F**).

**Figure 2.**
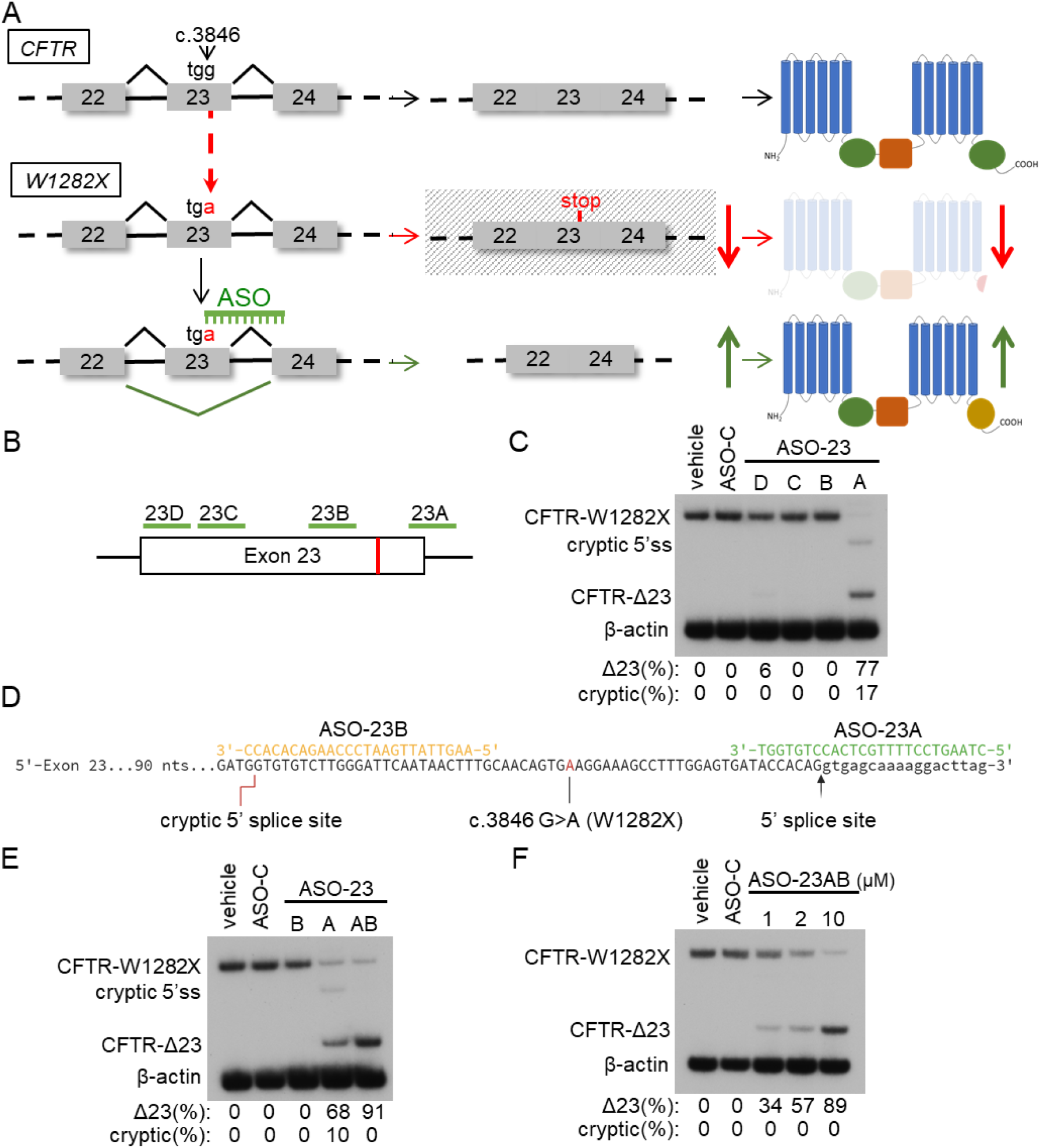
ASOs induce CFTR-W1282X exon 23 skipping. (**A**) Schematic of the CFTR dysfunction caused by the W1282X mutation in exon 23. The c.3846 G>A (W1282X) mutation creates a premature termination codon (PTC) in exon 23 leading to degradation of the transcript via NMD and reducing translation of a semi-functional truncated CFTR protein (indicated by red arrows). ASO induced exon 23 skipping, eliminates the PTC, restores CFTR mRNA stability and increases CFTR expression (green arrows). (**B**) Diagram of ASO target sites (green lines) on human CFTR exon 23 pre-mRNA. The location of the CFTR-W1282X mutation is indicated by a red vertical line. **(C)** RT-PCR analysis of CFTR exon 23 splicing in a CFF16HBEge-W1282X cell line treated with the indicated ASO (10μM). Exon 23 skipping or cryptic splice site activation was quantified [Δ23 or cryptic/(Δ23+cryptic+W1282X)x100] and is indicated below each lane. (**D**) Sequence alignment of ASO-23A and ASO-23B to exon 23. (**E**) RT-PCR analysis of exon 23 splicing in CFF16HBEge-W1282X cells treated with ASO-23A, ASO-23B, ASO-23A + ASO-23B (ASO-23AB, 10μM each), or ASO-C (20μM). Exon 23 skipping or cryptic splice site activation (% of total) was quantified and is indicated below each lane. (**F**) RT-PCR analysis of exon 23 splicing in CFF16HBEge-W1282X cells treated with ASO-23AB, or ASO-C at indicated concentrations. Exon 23 skipping and cryptic splice site activation was quantified and is indicated below each lane. β-actin is a control for RNA expression in all experiments.

We next tested whether ASO-23AB treatment could increase conductance in the immortalized hBE *CFTR-W1282X* cell lines. ASO-23AB treatment resulted in a significant increase in conductance when modulators were present compared to controls (**Figure 3A,B**). The conductance increased in an ASO dose-dependent manner (**Figure 3B**). This increase in activity corresponded with an increase in exon 23 skipping (**Figure 3C,D**). There was a positive correlation between ASO-induced exon 23 skipping and conductance (**Figure 3E**).

**Figure 3.**
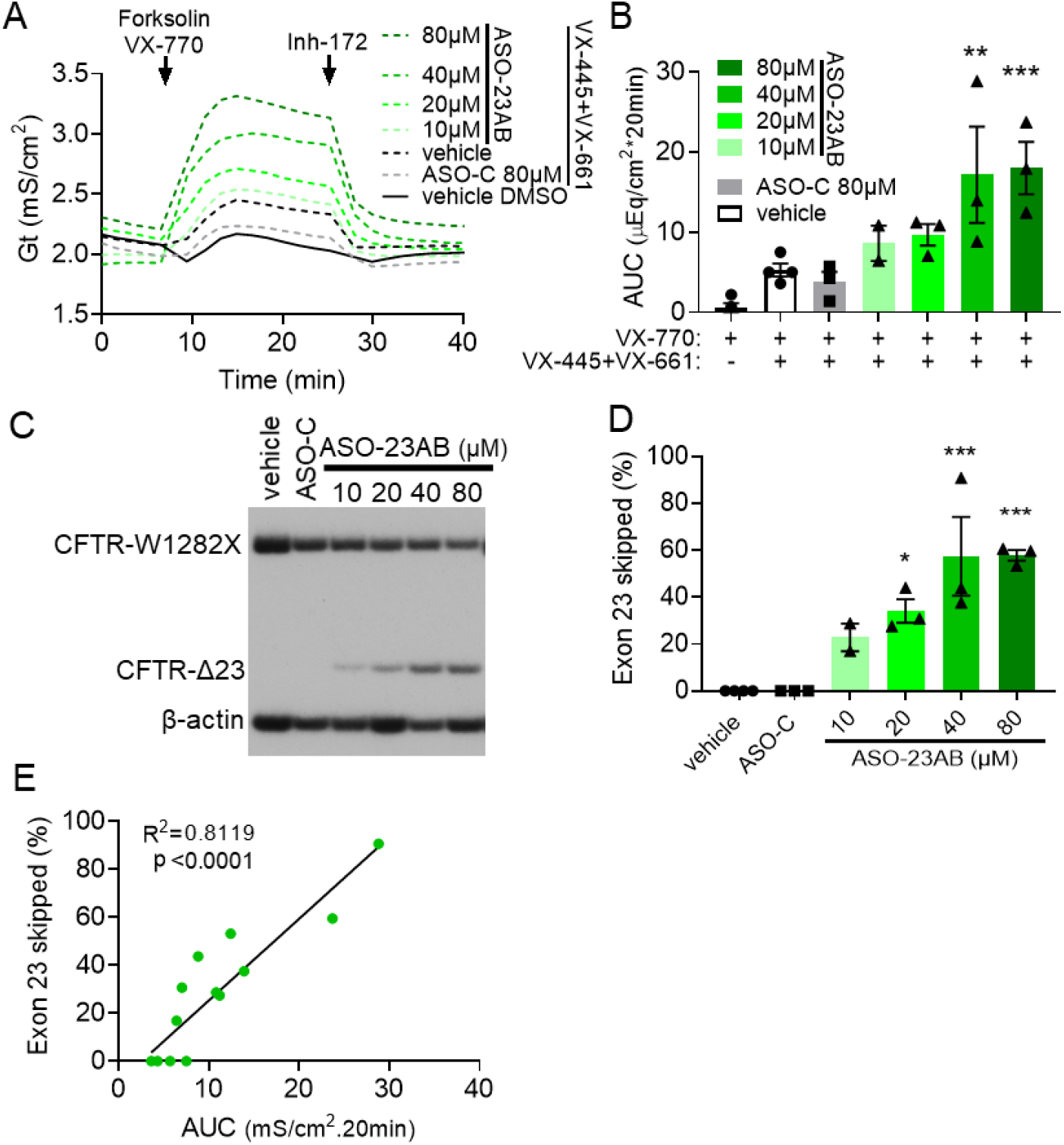
ASO-induced exon skipping increases membrane conductance of a CFTR-W1282X cell line. (**A**) Conductance (Gt) traces of a CFF16HBEge-W1282X clonal cell line selected for high resistance (CFF16HBEge-W1282X-SCC:3F2). Cells were transfected with vehicle (black line), ASO-C (grey line), or ASO-23AB (green lines) at indicated concentrations. Cells were pre-treated with DMSO (solid line) or VX-445+VX-661 (dashed lines). (**B**) Average area under the curve (AUC) of the conductance trace (A) was quantified for the forskolin + VX-770 test period for each treatment group. Error bars are ±SEM. One-way ANOVA; Dunnett’s multiple comparison test to vehicle + DMSO, **p<0.01, ***p<0.001. N=3 except for 10 μM ASO-23AB where N=2. (**C**) RT-PCR analysis of exon 23 splicing in CFF16HBEge-W1282X cells in (A). β-actin is a control for RNA expression. **(D)** Quantification of exon 23 skipping (% of total) in (C). Error bars are ±SEM. One-way ANOVA; Dunnett’s multiple comparison test to vehicle, *p<0.05, ***p<0.001. N=3 except for 10 μM ASO-23AB where N=2. (**E**) The calculated area under the curve, shown in (B) correlated with exon 23 skipping (%), shown in (D) (simple linear regression).

### ASO-induced exon skipping rescues chloride currents in homozygous *CFTR-W1282X* patient-derived bronchial epithelial cells

To further assess the therapeutic potential of ASO-induced exon skipping in correcting the *CFTR-W1282X* mutation, we analyzed the effects of ASO treatment on channel activity in differentiated primary human bronchial epithelial (hBE) cells isolated from a CF patient homozygous for *CFTR-W1282X*. This cell-based model is the gold-standard for pre-clinical testing of CF therapeutics as the functional responses to drugs in this assay has been shown to accurately predict efficacy in the clinic (35, 36). Transepithelial voltage (Vt) and resistance (Rt) was recorded to calculate an equivalent current (Ieq=Vt/Rt). Without ASO treatment, only the combination treatment of VX-770, VX-445 and VX-661 had a significant effect on chloride secretion in the cells (**Figure 4A,B**). ASO-23AB treatment in combination with VX-770 + C18, or VX-770 + VX-445 + VX-661, resulted in a ∼5-fold and 3-fold increase in chloride secretion, respectively, compared to either modulator treatment alone (**Figure 4A,B**).

**Figure 4.**
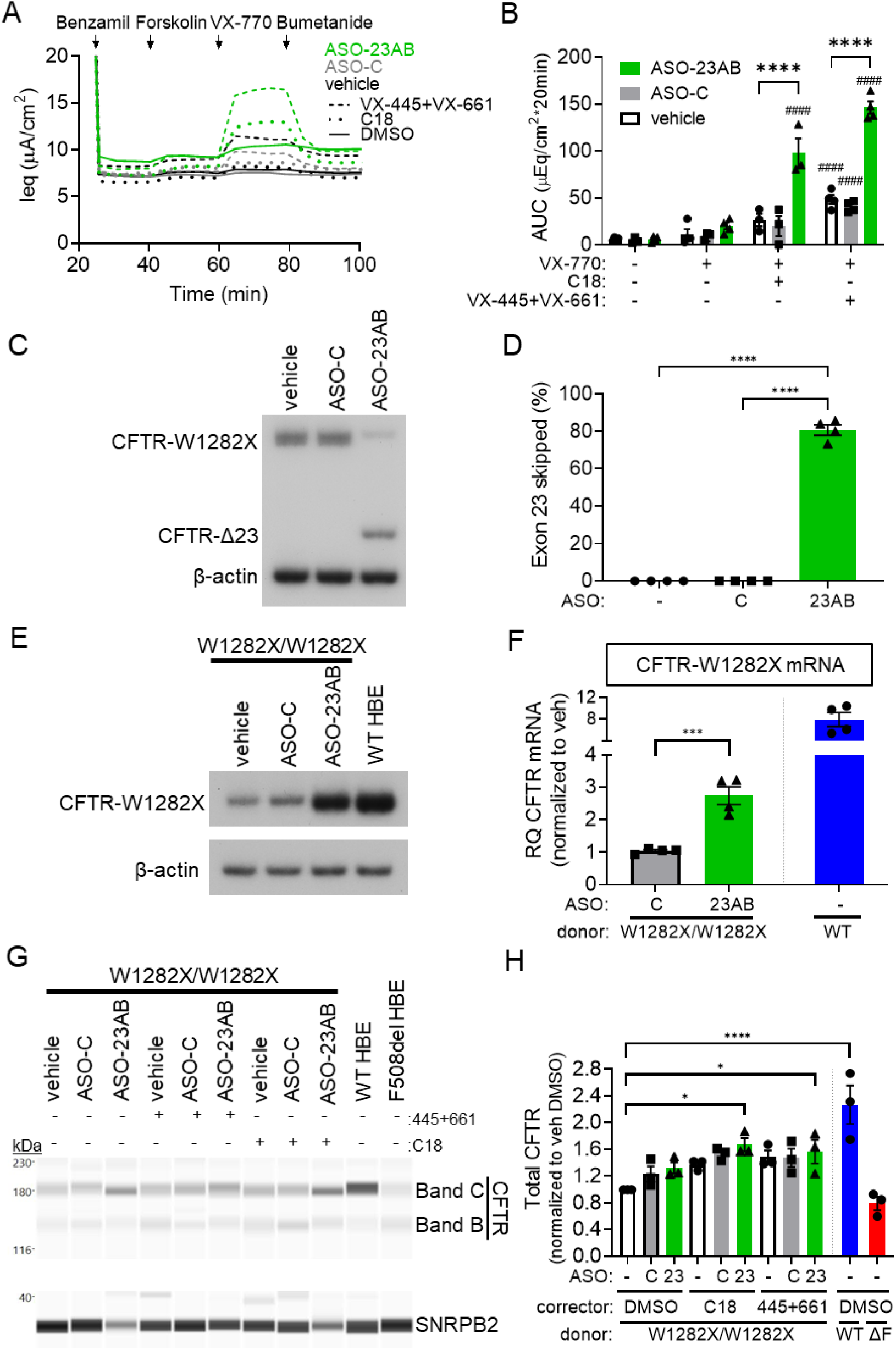
ASO treatment induces exon 23 skipping, stabilizes CFTR mRNA, and rescues CFTR function in primary human bronchial epithelial (hBE) cells isolated from a patient homozygous for *CFTR-W1282X*. (**A**) Equivalent current (Ieq) traces of primary hBE cells from a homozygous *CFTR-W1282X* CF donor. Cells were transfected with vehicle, ASO-C, or ASO-23AB (320 μM total) and pre-treated with DMSO, C18, or VX-445+VX-661. (**B**) Average AUC of the current traces (A) was quantified for the forskolin or forskolin + VX-770 test periods for each treatment group. Error bars are ±SEM. Two-way ANOVA; Dunnett’s multiple comparison test to DMSO within treatment groups, ####p<0.01. Two-way ANOVA; Dunnett’s multiple comparison test to vehicle within treatment groups, ****p<0.0001. N=4 except C18 treatment where N=3. (**C**) RT-PCR analysis of exon 23 splicing in cells from (A). β-actin is a control for RNA expression. (**D**) Quantification of exon 23 skipping in (C). Error bars are ±SEM. One-way ANOVA; Dunnett’s multiple comparison test, ****p<0.0001. N=4. (**E**) RT-PCR analysis of CFTR mRNA expression (exons 11-14) from cells analyzed in (A) compared to hBE cells from a non-CF donor. (**F**) RT-qPCR analyses of total CFTR mRNA (exon 11-12) from cells analyzed in (A) compared to hBE cells from a non-CF donor. Error bars are ±SEM. One-way ANOVA; Tukey’s multiple comparison test, ****p<0.0001. N=4. (**G**) Immunoblot analysis of CFTR protein isolated from cells in (A). Protein from a non-CF donor and a CF donor homozygous for F508del is also shown. SNRPB2 is a loading control. (**H**) Quantification of the Total CFTR (B+C Bands)/SNRP2 normalized to vehicle + DMSO shown in (G). Error bars are ±SEM. One-way ANOVA; Dunnett’s multiple comparison test to vehicle + DMSO, *p<0.05, ****p<0.0001. N=3.

This functional rescue by ASO-23AB treatment was accompanied by a significant induction of exon 23 skipping (**Figure 4C,D**). ASO-23AB treatment resulted in a 3-fold increase in total *CFTR* mRNA compared to untreated samples, a level that is ∼30% of mRNA levels in wildtype non-CF donor hBE cells (**Figure 4E,F**). The rescue of total *CFTR* RNA expression is indicative of a stabilization of mRNA as a result of elimination of the PTC introduced by the *CFTR-W1282X* mutation in exon 23. ASO treatment also resulted in an increase in CFTR protein (**Figure 4G,H**). The stabilization of *CFTR-W1282X* mRNA and increase in CFTR protein expression correlates with the rescue of chloride secretion in these patient cells, predictive of a potential therapeutic effect of ASO-23AB treatment over current modulator drugs for patients homozygous for *CFTR-W1282X*.

### Activity and allele-specificity of ASOs in patient-derived bronchial epithelial cells compound heterozygous for *CFTR-W1282X* and *F508del*

Many CF patients with the *CFTR-W1282X* mutation are compound heterozygotes, with a different mutation, most commonly *CFTR-F508del*, in the other *CFTR* allele. This second allele would also be a target of ASO-induced exon 23 skipping and would result in a CFTR protein with the original mutation and a deletion of exon 23. To test the effect of ASO-induced exon 23 skipping on mRNA from *CFTR* mutations commonly found with *CFTR-W1282X* in compound heterozygotes, we treated primary hBE cells isolated from a CF patient with the *CFTR-W1282X* and *CFTR-F508del* mutations, with ASO-23AB and measured chloride secretion (**Figure 5A**).

**Figure 5.**
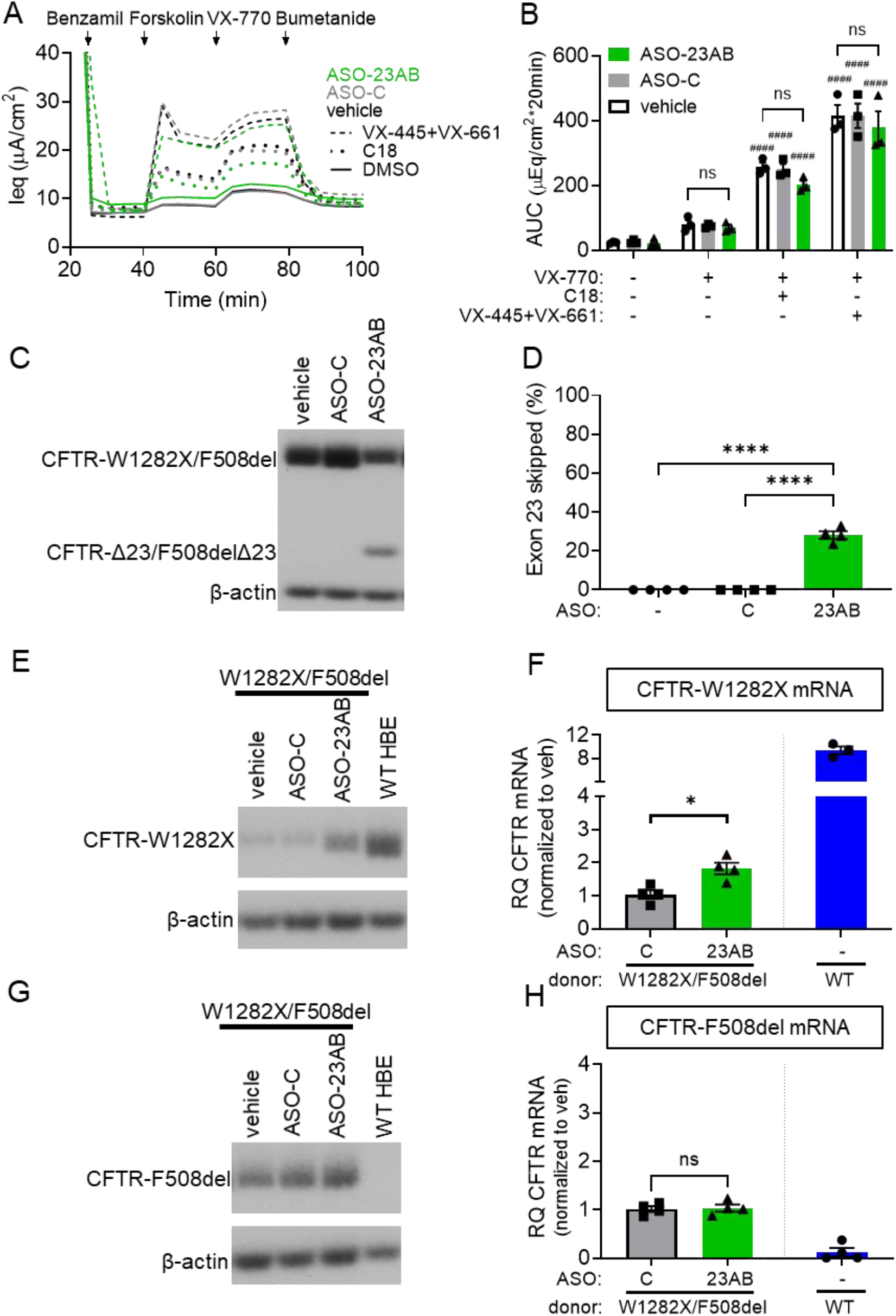
ASO treatment does not affect modulator activity in hBE cells isolated from a CF patient compound heterozygous for CFTR-W1282X and CFTR-F508del. (**A**) Equivalent current (Ieq) traces of primary hBE cells isolated from a CF donor heterozygous for CFTR-W1282X and CFTR-F508del. Cells were transfected with vehicle, ASO-C, or ASO-23AB (320 μM total). Cells were pre-treated with DMSO, C18, or VX-445+VX-661. (**B**) Average AUC of the current traces (A) was quantified for the forskolin + VX-770 test periods for each treatment group. Error bars are ±SEM. Two-way ANOVA; Dunnet’s multiple comparison test to DMSO within treatment groups, #p<0.05, ####p<0.01. Two-way ANOVA; Dunnet’s multiple comparison test to vehicle within treatment groups, ns=p>0.05. N=3. (**C**) RT-PCR analysis of exon 23 splicing in hBE CFTR-W1282X/F508del cells in (A). β-actin is a control for RNA expression. (**D**) Quantification of exon 23 skipping in (C). Error bars are ±SEM. One-way ANOVA; Dunnet’s multiple comparison test, **p<0.01. N=3. (**E**) RT-PCR analysis of non-F508del CFTR mRNA expression (exons 11-14) from cells analyzed in (A) compared to hBE cells from a non-CF donor. (**F**) RT-qPCR analyses of total CFTR mRNA from non-F508del alleles (exon 11-12) in cells analyzed in (A) compared to hBE cells from a non-CF donor. Error bars are ±SEM. One-way ANOVA; Tukey’s multiple comparison test, ****p<0.0001. N=4, one outlier identified and removed with ROUT outlier analysis (Q=5%) in WT. (**G**) RT-PCR analysis of F508del CFTR mRNA expression (exons 11-14) from cells analyzed in (A) compared to hBE cells from a non-CF donor. (**H**) RT-qPCR analyses of total CFTR mRNA from F508del alleles (exon 11-12) in cells analyzed in (A) compared to hBE cells from a non-CF donor. Error bars are ±SEM. One-way ANOVA; Tukey’s multiple comparison test, ns=p>0.05. N=4.

In cells from this patient, both modulator combinations of VX-770 + C18, and VX-770 + VX-445 + VX-661, resulted in significant recovery of chloride secretion, as expected given that *CFTR-F508del* is known to be responsive to each drug (**Figure 5B**). ASO-23AB treatment had no significant effect on this rescue, with cells showing no increase or decrease in potentiator and corrector response (**Figure 5B**).

Analysis of RNA splicing revealed a significant induction of exon 23 skipping with ASO treatment compared to the controls. However, exon 23 skipping was considerably lower (25% of total RNA) (**Figure 5C,D**) than the skipping obtained in *CFTR-W1282X* homozygous donor cells (80% of total RNA) (**Figure 4C,D**). These results suggest that exon 23 skipping of RNA from *CFTR-F508del* may be less efficient.

To analyze the effect of ASO-23AB on exon 23 from mRNA derived from each allele specifically, primers were designed to anneal at the F508del mutation site and specifically amplify either non-F508del or F508del mRNA (**Table S1**). When comparing the baseline expression of mRNA from each allele, RNA derived from the *CFTR-W1282X* allele was only 20% of that expressed from the *CFTR-F508del* allele (**Figure S1A**) most likely due to transcript degradation by nonsense mediated decay. ASO treatment significantly increased this expression to ∼40% of that generated from the *CFTR-F508del* allele (**Figure S1A**). Analysis of RNA from each allele separately revealed a 2-fold increase in *CFTR* transcripts from *CFTR-W1282X*, similar to levels achieved in the homozygous donor and up to 20% of wild-type *CFTR* expression (**Figure 5E,F**). In contrast, the ASO had no significant effect on total *CFTR* mRNA expression from the *CFTR-F508del* allele (**Figure 5G,H**).

To further investigate the reason for an increase in total mRNA from the *CFTR-W1282X* allele with ASO treatment, yet a large reduction in total exon 23 skipping compared to the homozygous donor cells, we analyzed ASO-induced exon 23 skipping from each allele using allele specific primers. This analysis revealed an increase in exon skipping in mRNA from the *CFTR-W1282X* allele (71%) compared to transcripts from the *CFTR-F508del* allele (2%), suggesting that ASO treatment has a greater effect on *CFTR-W1282X* than on *CFTR-F508del* (**Figure S1B**). Comparative sequence analysis of exon 23 indicated that binding sites for several splicing proteins are eliminated by the G>A change in W1282X compared to wild-type CFTR (ESEfinder3.0) (**Figure S1C**) (37). The elimination of these splicing enhancer cis-acting sequences may weaken splicing to the exon and make the ASO more effective in inducing skipping. In fact, ASO treatment in non-CF and *CFTR-F508del* homozygous donor cells in comparison to donor cells with either one or two copies of *CFTR-W1282X* showed an increase in ASO-induced exon 23 skipping that correlated with the number of *CFTR-W1282X* alleles (**Figure S1D**). This result suggests ASO-23AB may be more effective at inducing exon 23 skipping in *CFTR-W1282X* RNA, which could be advantageous in treating CF patients compound heterozygous for *CFTR-W1282X* and another *CFTR* mutation less responsive to modulator treatment.

## Discussion

Despite clinical success, the use of ASOs to correct the translational open reading frame and recover gene expression in diseases caused by frameshift or nonsense mutations resulting in PTCs has not been extensively explored as a therapeutic approach. These types of mutations are the most common disease-causing mutations and account for ∼20% of disease-associated mutations in cystic fibrosis. Here we demonstrate that elimination of the relatively common nonsense mutation, *CFTR-W1282X*, by ASO-induced skipping of *CFTR* exon 23, which encodes the mutation, recovers CFTR expression. The activity of this CFTR isoform, lacking 52 amino acids, requires CFTR modulator drugs that are currently used to treat CF patients. Thus, this ASO approach in combination with current CF drugs offers a potential therapeutic for individuals with the *CFTR-W1282X* mutation and opens the door for similar strategies to treat other terminating mutations, both in CF and other diseases.

Gene mutations that result in PTCs are challenging to treat because they not only encode a truncated protein product, but the mRNA intermediate is a target of NMD, a cellular quality-control mechanism whereby mRNA with PTCs are degraded (38, 39). Studies have shown that, if produced at sufficient levels, CFTR-W1282X protein is responsive to current CFTR modulator therapies (16, 18, 19, 21, 22) but because stop mutations in CFTR result in a dramatic loss of *CFTR* expression, CF patients are not usually responsive to these drugs. Approaches are being pursued to identify molecules that stabilize mRNA by blocking NMD but, because NMD is an important mechanism regulating gene expression, any approach must avoid global inhibition of NMD which would likely have toxic effects (21, 23). Small molecules that promote translational readthrough of termination codons are also being explored as potential treatments for PTC mutations, including CF (40–45). However, effects from the long-term use of these drugs has raised concerns (44, 46, 47). Though these approaches may hold promise, none are specific to *CFTR* directly, and to date none have been approved for use in CF patients. More recently, gene-specific suppression of NMD, has been explored as a promising approach to overcome potential risk of global NMD knockdown (48, 49).

Using ASOs to remove exons encoding stop codons to correct the translational open reading frame has broad applications for addressing terminating mutations. ASO-mediated reading frame correction via induced skipping of symmetrical exons has shown promise in the FDA-approved ASOs targeting PTCs in Duchenne’s Muscular Dystrophy (50) and also in pre-clinical studies in mice for the treatment of diseases such as CLN3 Batten (51). A critical requirement for this approach is that the induced protein isoform must retain partial function. We show that CFTR-Δ23 had significant cAMP-activated conductance responses that were further enhanced by modulator treatments (**Figure 1**). Exon 23 encodes amino acids near the C-terminus of CFTR including a portion of the nucleotide-binding domain 2 (NBD2). The retained function of CFTR-Δ23 is consistent with previous reports that truncation at NBD2 results in a CFTR protein that is trafficked to the cell surface, albeit with deleterious effects on channel gating (26, 28). The result also aligns with data showing that the *CFTR-W1282X* mutation is responsive to modulator therapies (16–18) (**Figure 1**). Notably, though truncation of CFTR at NBD2 results in some retained function, domains at the C-terminus are important for stability and gating and these domains are preserved when exon 23 is skipped (27, 28, 52, 53).

The clinical potential of ASO delivery to the respiratory system, one of the primary targets for CF therapeutics, has been demonstrated for asthma and other inflammatory lung conditions (54–57). Naked ASOs have been successfully delivered to the lung, where they access multiple cell types including cells which express *CFTR* (55, 58–60). Aerosolization of ASOs have shown promise in delivery in both a CF-like lung disease model in mice as well as CF patients (59, 61, 62). Additionally, ASOs have been shown to have long-lasting effects in primary human bronchial epithelial cells isolated from CF-patients (63), and *in vivo*, with treatment durations lasting weeks to months before further dosing is needed in patients (24, 64–66). In addition, as patients with class I mutations tend to have disease phenotypes in other organs including the digestive system, systemic delivery of ASOs may also be considered (67, 68). Intravenous and subcutaneous injections of ASOs are currently used to treat patients and bioavailable ASO formulations that target the gut epithelia have shown some potential (69–71).

Unlike other therapeutics targeting global NMD or inducing translational readthrough, our ASO targets *CFTR* transcripts specifically to induce skipping of exon 23 (**Figure 2**). ASO treatment in immortalized and patient-derived hBE cells expressing *CFTR-W1282X* resulted in a dose-dependent increase in exon 23 skipping that correlated with an increase in function (**Figure 3,4**). Further, assessment of mRNA from ASO-treated primary hBE cells from a patient with *CFTR-W1282X* and *CFTR-F508del* revealed partial allele specificity of ASO induced exon 23 skipping for CFTR-W1282X (**Figure S1**), potentially broadening the scope of this ASO strategy to CF patients heterozygous for *CFTR-W1282X* and another CFTR mutation less responsive to current modulator therapies. Despite effectively inducing skipping of *CFTR-W1282X* mRNA, ASO-induced exon 23 skipping did not have an improved effect on function compared to modulator treatment alone (**Figure 5**). It is possible that a functional ceiling is achieved with modulator treatment in cells from this compound heterozygous donor, and additional expression from the *CFTR-W1282X* allele does not increase chloride current above what was already achieved with modulator rescue *on CFTR-F508del*, for which the modulators were developed. There have been a number of studies identifying other corrector and potentiator drugs superior to these modulators in the context of *CFTR-W1282X* (16, 19, 21, 72). The effect of these new modulators was not tested along with ASO treatment here, but they have been shown to be effective in rescuing *CFTR-W1282X* activity in conjunction with other readthrough compounds that enhance *CFTR-W1282X* expression (21, 72). Future studies may reveal better modulator/ASO combinations for treating *CFTR-W1282X* compound heterozygotes. Overall, our results support the use of ASO treatment in combination with approved CF modulators as an effective treatment option for CF patients with class I mutations within symmetrical exons.

## Materials and Methods

### Expression plasmids

CFTR-Δ23 and CFTR-W1282X were created from the synthetic *CFTR* high codon adaption index (HCAI) construct subcloned in the pcDNA3.1/Neo(+) vector (73) using the Q5 Site-Directed Mutagenesis Kit (NEB) with primers flanking each exon (Table S1). All plasmids were sequenced to confirm mutations. Plasmids were stably transfected into Fischer Rat Thyroid (FRT) cells in 6-well plates using lipofectamine LTX (Thermo Fisher) and OptiMEM (Thermo Fisher) for 48 hours. Cells were transferred to T75 flasks and clonal cell lines were selected with G418 (300 μg/ml) for one week. After selection cells were maintained in media supplemented with G418 (150 μg/ml).

### Cells and culture conditions

FRT cell lines were cultured in F12 Coon’s modification media (Sigma, F6636) supplemented with 10% FBS and 1% Penicillin-Streptomycin (PenStrep). 16hBEge-W1282X cell lines were obtained from the Cystic Fibrosis Foundation (CFF) and cultured according to their instructions in MEM media (34, 74). Single-cell clones of the original CFF16hBEge-W1282X cell line were created to select for high resistance clonal cell lines. One clonal cell line, CFF16hBEge-W1282X-SCC:3F2, was selected for analysis. Primary human bronchial epithelial cells (hBE) isolated from CF patients homozygous for *CFTR-W1282X* (patient code HBEU10014) and compound heterozygous for *CFTR-W1282X* and *CFTR-F508del* (patient code HBEND12112) were also obtained from the CFF. For functional analysis cells were differentiated by plating on Costar 24-well high-throughput screening filter plates (0.4 μM pore size, Polyester, Corning, catalog #CLS3397). FRT and 16hBE cells were grown in a liquid/liquid interface (180 μl apical/700 μl basolateral) in a 37°C incubator with 90% humidity and 5% CO_2_ for one week. Primary hBE cells were differentiated in an air/liquid interface for five weeks. Media was replaced three times a week.

### Antisense oligonucleotides

Splice-switching antisense oligonucleotides are 25-mer phosphorodiamidate morpholino oligomers (Gene-Tools, LLC) (Table S1). A non-targeting PMO was used as a negative control, ASO-C, (Gene Tools, standard control oligo). ASOs were formulated in sterile water.

### ASO cell transfection

For splicing analysis CFF16hBEge-W1282X-SCC:3F2 clones were transfected with ASOs at indicated concentrations on 24-well plates in Minimum Essential Medium (MEM) media supplemented with 10% FBS and 1% PenStrep. Cells were transfected using Endo-Porter (Gene-Tools, 6 μl/ml) for 48 hours (75).

For functional analysis CFF16HBEge-W1282X-SCC:3F2 cells were transfected on filter plates four days post plating. Cells were transfected with ASOs apically in 100 μl of complete MEM media with Endo-Porter at indicated concentrations for 48 hours.

Primary hBE cells were transfected after differentiation on filter plates as previously described (63). Briefly, cells were transfected with ASO in an apical hypo-osmotic solution for 1 hour. The solution was removed, and the cells were treated again in DPBS for 4 days until functional analysis.

### RNA isolation and RT-PCR

RNA was extracted from cells using TRIzol according to manufacturer instructions (Thermo Fisher Scientific). Reverse transcription was performed on total RNA using the GoScript Reverse Transcription System with an oligo-dT primer (Promega). Splicing was analyzed by radiolabeled PCR of resulting cDNA using GoTaq Green (Promega) spiked with α-^32^P-deoxycytidine triphosphate (dCTP). Primers for amplification are reported in Table S1. Reaction products were run on a 6% non-denaturing polyacrylamide gel and quantified using a Typhoon 7000 phosphorimager (GE Healthcare) and ImageJ software (76).

### Real-time qPCR

Real-time qPCR was performed with PrimeTime Gene Expression Master Mix and PrimeTime qPCR probe assay kits human non-F508del-CFTR (IDT, qhCFTR-ex11WTF, qhCFTR-ex12WTR, hCFTR-F508), and human F508del-CFTR (IDT, qhCFTR-ex11ΔFF, qhCFTR-ex12ΔFR, hCFTR-DF508) transcripts were normalized to human HPRT1 (IDT, Hs.PT.58v.45621572) (**Table S1**). All reactions were analyzed in triplicate on 96-well plates and averaged together to comprise one replicate. Real-time PCR was performed on an Applied Biosystems (ABI) ViiA 7 Real-Time PCR System.. Results were analyzed by the ΔΔCT method (77).

### Protein isolation and automated western analysis

Cell lysates for immunoblot analysis were prepared from cells after functional analysis using NP-40 lysis buffer (1% Igepal, 150mM NaCl, 50mM Tris-HCl pH7.6) supplemented with 1× protease inhibitor cocktail (Sigma-Aldrich, cat #11836170001). Protein concentration was measured using a Coomassie (Bradford) protein assay (Thermo Fisher, cat #23200). Cell lysates were prepared using the sample preparation kit (Protein Simple) for an automated capillary western blot system, WES System (Protein Simple) (78, 79). Cell lysates were mixed with 0.1× sample buffer and 5× fluorescent master mix for a final protein lysate concentration of 0.2 mg/ml (FRTs) or 1.5 mg/ml (hBEs). Samples were incubated at room temperature for 20 minutes and then combined with biotinylated protein size markers, primary antibodies against CFTR 432 (FRTs), 570 + 450 (hBEs) (Riordan lab UNC, Cystic Fibrosis Foundation, diluted 1:100, or 1:50 + 1:200, with milk-free antibody diluent), β-actin (C4, Santa Cruz Biotechnology, diluted 1:50 with milk-free antibody diluent), and SNRBP2 (4G3, provided by the Krainer Lab; B’’, diluted 1:2000 with milk-free antibody diluent), horseradish peroxidase (HRP)-conjugated secondary antibodies, chemiluminescence substrate and wash buffer and dispensed into respective wells of the assay plate and placed in WES apparatus. Samples were run in duplicate or triplicate. Signal intensity (area) of the protein was normalized to the peak area of the loading control C4, β-actin (FRT) or B’’, SNRPB2 (hBE). Quantitative analysis of the CFTR B and C-bands was performed using Compass software (Protein Simple).

### Automated conductance and equivalent current assay

Stably transfected FRT cells, 16HBEge-W1282X-SCC:3F2, and primary hBE cells were treated with C18 (6 μM) (VRT-534, VX-809 analog), VX-445 + VX-661 (3 μM + 3.5 μM FRT, 1 μM + 3 μM 16hBE and primary hBE) or vehicle (equivalent DMSO) at 37°C (33). Twenty-four hours later the cells were switched from differentiation media to HEPES-buffered (pH 7.4) F12 Coon’s modification media (Sigma, F6636) apically and basolaterally and allowed to equilibrate for one hour at 37°C without CO_2_. To obtain the conductance measurements, the transepithelial resistance was recorded at 37°C with a 24-channel TECC robotic system (EP Design, Belgium) as previously described (80). Briefly, for the FRT cells, baseline measurements were taken for ∼20 minutes. Forskolin (10 μM) was first added to the apical and basolateral sides and then cells were treated with potentiator, VX-770 (1 μM). Finally, inhibitor-172 (Inh-172, 20 μM) was added to inactivate CFTR. The 16hBEge-W1812X-3F2 clones were measured similarly apart from forskolin and VX-770 added to the cells at the same time. Measurements were taken at two-minute intervals. Gt was calculated by the reciprocal of the recorded Rt (Gt=1/Rt), after Rt was corrected for solution resistance (Rs) and plotted as conductance traces (**Figures 1C,3A**). Calculated equivalent currents (I_eq_) were obtained similar and as outlined in Michaels et al (63). I_eq_ was calculated using Ohm’s law (I_eq_ = Vt/Rt) and plotted as current traces (**Figures 4A,5A**). To estimate average functional response trajectories during each test period, area under the curve measurements of forskolin and forskolin + VX-770 were calculated using a one-third trapezoidal rule for each test period using Excel. The average of two identically treated wells was calculated for each plate to obtain one biological replicate used in the final mean ±SEM graphed.

### Statistics

Statistical analyses were performed using GraphPad PRISM 9.2.0. The specific statistical test used in each experiment can be found in the figure legends.

## Supplemental Material

Supplemental material can be found online.

## Acknowledgments

We thank the Cystic Fibrosis Foundation for providing the immortalized and patient hBE cells. We thank Adrian Krainer, Cold Spring Harbor Laboratory, for use of the 4G3 antibody. We also thank the Hastings and Bridges labs for assistance with experiments and insightful discussions, in particular Francine Jodelka and Dr. Jessica Centa, for their help in editing the manuscript. This work was partially funded by a grant from the Cystic Fibrosis Foundation.

