## Supplementary material for "Open reading frame correction using antisense oligonucleotides for the treatment of cystic fibrosis caused by CFTR-W1282X": Figure S1, Table S1

**Supplemental Material**

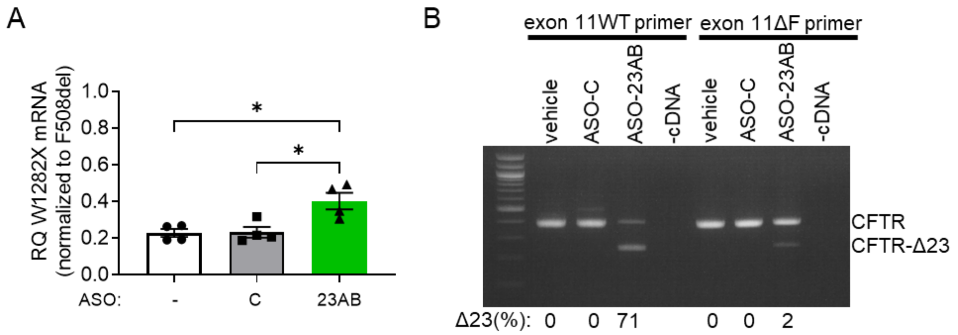


C

CFTR exon 23:
GTGGGCCTCTTGGGAAGAACTGGATCAGGGAAGAGTACTTTGTTATCAGCTTTTTTGAGACTACTGAACACTGAAGGA
GAAATCCAGATCGATGGTGTGTCTTGGGATTCAATAACTTTGCAACAGTGGAGGAAAGCCTTTGGAGTGATACCACAG

| SRSF1 threshold: 1.956 | SRSF1 (IgM-BRCA1) threshold: 1.867 | SRSF2 threshold: 2.383 | SRSF5 threshold: 2.67 | SRSF6 threshold: 2.676 |
|---|---|---|---|---|
| Position/Site/Score | Position/Site/Score | Position/Site/Score | Position/Site/Score | Position/Site/Score |
| 57 GAGACTA 2.64552 | 71 CTGAAGG 2.2387 | 4 GGCCTCTT 3.43225 | 58 AGACTAC 2.79705 | 96 TGTGTC 4.58418 |
| 71 CTGAAGG 1.96982 | 124 CAGTGGA 3.15115 | 5 GCCTCTTG 2.45572 | 61 CTACTGA 3.02819 | 119 TGCAAC 2.92201 |
| 124 CAGTGGA 3.28787 | 127 TGGAGGA 2.00488 | 59 GACTACTG 5.38755 | 123 ACAGTGG 3.30474 | |
| 127 TGGAGGA 2.34103 | | 66 GAACACTG 4.60771 | | |
| | | 87 GATCGATG 3.3474 | | |
| | | 105 GGATTCAA 2.97789 | | |
| | | 149 TACCACAG 2.54564 | | |

CFTR W1282X:
GTGGGCCTCTTGGGAAGAACTGGATCAGGGAAGAGTACTTTGTTATCAGCTTTTTTGAGACTACTGAACACTGAAGGA
GAAATCCAGATCGATGGTGTGTCTTGGGATTCAATAACTTTGCAACAGTGAAGGAAAGCCTTTGGAGTGATACCACAG

| SRSF1 threshold: 1.956 | SRSF1 (IgM-BRCA1) threshold: 1.867 | SRSF2 threshold: 2.383 | SRSF5 threshold: 2.67 | SRSF6 threshold: 2.676 |
|---|---|---|---|---|
| Position/Site/Score | Position/Site/Score | Position/Site/Score | Position/Site/Score | Position/Site/Score |
| 57 GAGACTA 2.64552 | 71 CTGAAGG 2.2387 | 4 GGCCTCTT 3.43225 | 58 AGACTAC 2.79705 | 96 TGTGTC 4.58418 |
| 71 CTGAAGG 1.96982 | | 5 GCCTCTTG 2.45572 | 61 CTACTGA 3.02819 | 119 TGCAAC 2.92201 |
| | | 59 GACTACTG 5.38755 | | |
| | | 66 GAACACTG 4.60771 | | |
| | | 87 GATCGATG 3.3474 | | |
| | | 105 GGATTCAA 2.97789 | | |
| | | 149 TACCACAG 2.54564 | | |

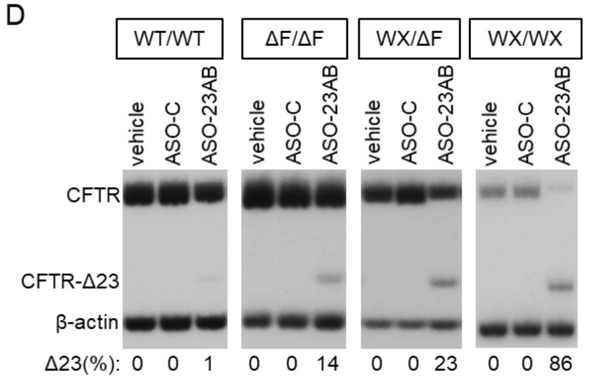


**Figure S1.** ASO-induced exon 23 exclusion has partial allele specificity for CFTR-W1282X (**A**) RT-qPCR analyses of allele-specific mRNA expression isolated from compound heterozygous hBE cells shown in Figure 5 F&H. Total CFTR mRNA expression from the CFTR-W1282X allele was normalized to mRNA expression from the F508del allele for each treatment group. Error bars are ±SEM. One-way ANOVA; Tukey's multiple comparison test, *p<0.05. N=2. (**B**) Allele specific RT-PCR analysis of ASO-induced exon 23 skipping in hBE cells from a donor heterozygous shown in Figure 5A. Allele specific transcripts were amplified from exons 11-25 (primers: 11WT-25 [W1282X] or 11ΔF-25 [F508del]). Exon 23 skipping from each allele was analyzed using nested exon 22-24 primers. Exon 23 skipping was quantified (% of total) and is shown below each lane. (**C**) Comparison of calculated SR protein binding sites (ESEfinder, Cold Spring Harbor Laboratory, (37)) between WT exon 23 (top) and exon 23 containing the CFTR-W1282X mutation (bottom). Differences are indicated in red. (**D**) RT-PCR analysis of CFTR exon 23 skipping in hBE cells from various donors treated with vehicle, ASO-C, or ASO-23AB (320 μM). Genotype of each donor is indicated. Exon 23 skipping was quantified and is indicated below each lane. β-actin is a control for RNA expression.

**Table S1.** Splice-switching antisense oligonucleotide, primers, and probes used in the manuscript

| ASOs | Sequence (5'-3') |
|---|---|
| ASO-23A | CTAAGTCCTTTTGCTCACCTGTGGT |
| ASO-23B | AAGTTATTGAATCCCAAGACACACC |
| ASO-23C | AGCTGATAACAAAGTACTCTTCCCT |
| ASO-23D | ATCCAGTTCTTCCCAAGAGGCCCAC |
| ASO-C | CCTCTTACCTCAGTTACAATTTATA |
| **Primers** | **Sequence (5'-3')** |
| 1:HCAI-CFTRdel23R | GCGCTGGCCGGGGCTGAT |
| 2:HCAI-CFTRdel23F | AAGGTGTTCATCTTCAGCGGCACCTTC |
| 3: HCAI-CFTRWXF | TGCAGCAGTGACGCAAGGCCT |
| 4: HCAI-CFTRWXR | GGGTGATGCTGTCCCAGC |
| 5: hCFTR-ex11ΔFF | GCCTGGCACCATTAAAGAAAATATCATTGG |
| 6: hCFTR-ex11F | GCCTGGCACCATTAAAGAAAATATCATCTT |
| 7: hCFTR-ex14R | TCCAGGAGACAGGAGCATCT |
| 8: hCFTR-ex22F | CCAAACCATACAAGAAT |
| 9: hCFTR-ex24R | GATCACTCCACTGTTCAT |
| 10: hCFTR-ex25R | GTTCTATCACAGATCTGAG |
| 11: qhCFTR-ex11WTF | TGGCACCATTAAAGAAAATATCATCTT |
| 12: qhCFTR-ex12WTR | CTCAGTGTGATTCCACCTTCTC |
| 13: qhCFTR-ex11ΔFF | GGCACCATTAAAGAAAATATCATTGG |
| 14: qhCFTR-ex12ΔFR | CTCAGTGTGATTCCACCTTCT |
| 15: hβ-actinFor | AAAGACCTGTACGCCAACAC |
| 16: hβ-actinRev | GTCATACTCCTGCTTGCTGAT |
| 17: qhHPRT1For | GCGATGTCAATAGGACTCCAG |
| 18: qhHPRT1Rev | TTGTTGTAGGATATGCCCTTGA |
| **Probes** | **Sequence (5'-3')** |
| hCFTR-DF508 | /56-FAM/ACAGAAGCG/ZEN/TCATCAAAGCATGCC/3IABkFQ/ |
| hCFTR-F508 | /5.6-FAM/ACAGAAGCG/ZEN/TCATCAAAGCATGCC/3IABkFQ/ |
| hHPRT1 | /5HEX/AGCCTAAGA/ZEN/TGAGAGTTCAAGTTGAGTTTGG/3IABkFQ/ |
